# Learning progress mediates the link between cognitive effort and task engagement

**DOI:** 10.1101/2021.12.02.470970

**Authors:** Ceyda Sayalı, Emma Heling, Roshan Cools

## Abstract

While a substantial body of work has shown that cognitive effort is aversive and costly, a separate line of research on intrinsic motivation suggests that people spontaneously seek challenging tasks. According to one prominent account of intrinsic motivation, the Learning Progress Motivation theory, the preference for difficult tasks reflects the dynamic range that these tasks yield for changes in task performance (Oudeyer, Kaplan & Hafner, 2007). Here we test this hypothesis, by asking whether greater engagement with intermediately difficult tasks, indexed by subjective ratings and objective pupil measurements, is a function of trial-wise changes in performance. In a novel paradigm, we determined each individual’s capacity for task performance and used difficulty levels that are too low, intermediately challenging or high for that individual. We demonstrated that challenging tasks resulted in greater liking and engagement scores compared with easy tasks. Pupil size tracked objective task difficulty, where challenging tasks were associated with greater pupil responses than easy tasks. Most importantly, pupil responses were predicted by trial-to-trial changes in average accuracy as well as learning progress (derivative of average accuracy), while greater pupil responses also predicted greater subjective engagement scores. Together, these results substantiate the Learning Progress Motivation hypothesis stating that the link between task engagement and cognitive effort is mediated the dynamic range for changes in task performance.

## Introduction

According to the myth, Sisyphus was punished by the Gods to roll a boulder up a hill for eternity. It was the effort of his task that was considered a punishment for him. Although it is intuitive to assume that effort is aversive, people voluntarily engage throughout their lifespan in physically and mentally effortful tasks that allow them to acquire challenging hobbies, master expertise and be successful in adult life. Take video games for an example. Anyone who has played video games would agree that once they master a game level, they like to move onto a harder one. Most gamers would not voluntarily choose to play an easier level in order to avoid the effort. In fact, previous research has shown that when people are given the option, they gradually increase the difficulty of a video game across time (Baranes, Oudeyer & Gottlieb, 2014). In these games, participants keep their task accuracy around 50% on average, even if they can choose to play the easier task level the entire time. An important question, then, concerns what underlies the preference for specific effortful tasks while others are deemed frustrating. In line with the theories on intrinsic motivation (Gottlieb & Oudeyer, 2018) as well as the recent proposal (Agrawal et al., 2021) that the utility of mental tasks increase as a function of the information they provide, we test the hypothesis that effortful tasks are engaging if they yield an opportunity for increases in performance success.

Influential theories of effort assume that cognitive effort holds an intrinsic cost (Shenhav, Botvinick & Cohen, 2013). As such, cognitive effort discounting and selection paradigms (Westbrook, Kester & Braver, 2013) show that participants forego reward to avoid the performance of challenging tasks. However, a separate line of research on motivation suggests that humans can be intrinsically motivated for optimal challenge. For example, intrinsically motivating activities that are intermediately challenging given one’s capacity are known to induce a state of ‘flow’ (Csikszentmihalyi, 1990). In these activities, people report disliking tasks that are too easy or too difficult, but report greater engagement with and liking of those tasks for which accuracy is around 50%. Moreover, neural networks recruited during the experience of flow correspond to regions that are commonly associated with reward receipt (Ulrich, Keller & Grön, 2016).

One prominent account of intrinsic motivation is Learning Progress Motivation theory (Oudeyer, Kaplan & Hafner, 2007). This account suggests that the motivational value of a task reflects its potential for changes in task performance (Oudeyer, Gottlieb & Lopes, 2016).

Imagine three different types of task researchers might need to perform for work. Say, one is a task they know how to execute by heart, such as subject data entry. They know they will perform this job with almost perfect accuracy, and they will succeed in doing so. This means the changes in task performance for this task will be close to zero and will not change over time. Next, consider another task that is impossible for them to perform accurately, such as troubleshooting an fMRI machine. They know they will not be able to perform this task successfully, because they have no engineering background. So again, there will not be any changes in task performance. However, now consider a third job: a novel data analysis method that is intermediately challenging given their prior experience and associated with sufficient room for performance changes. They will fail once or twice and upon persistence, they will succeed, which will indicate to them that although difficult, the task is learnable, boosting the utility of this activity.

According to Learning Progress Motivation theory, these intermediately challenging tasks are exactly the tasks in which internally motivated agents must invest effort. Consistent with this framework, recent work showed that people are sensitive to both their average accuracy and their learning progress (Ten et al., 2021): People explored more difficult activities (associated with reduced accuracy) to the extent they were more learnable (providing a greater range of performance change). In a simulated experiment, researchers showed that artificial agents spontaneously spend more time exploring tasks that provide an opportunity for changes in task performance and avoid tasks that yield no change (Gottlieb & Oudeyer, 2018). Similarly, human infants have been shown to attend to intermediately predictable auditory and visual stimuli (Kidd, Piantadosi & Aslin, 2014; Kidd, Piantadosi & Aslin, 2012). Based on this account, Sisyphus’ punishment might have been reduced to the extent he experienced room for improving his performance in moving that boulder up the hill. Therefore, in the current study, we test the hypothesis that, relative to easy and difficult tasks, intermediately challenging tasks are perceived as more engaging, as indexed both by subjective report of engagement as well as by trial-by-trial objective indices of pupil dilation, which has been well established to be associated with task engagement (Aston-Jones & Cohen, 2005). Furthermore, based on the Learning Progress Motivation hypothesis, we anticipate that the trial-by-trial index of pupil dilation is predicted not just by average performance accuracy, but also by changes in performance accuracy.

To test these hypotheses, we leveraged a paradigm that is commonly used to induce a state of flow and engagement rather than avoidance of cognitive effort. After determining each individual’s maximum cognitive capacity, participants performed 4 blocks of effortful tasks in which they scored 25, 50, 75 and 100% correct. After performance of this second effort exposure phase, participants completed self-report questions about their subjective liking, perceived ability and engagement during these different task blocks. We predicted that intermediately challenging tasks would be perceived as more engaging than easy or difficult tasks, yielding an inverted-U relationship between task engagement and task accuracy. We also predicted that subjective engagement scores would vary both the average performance accuracy (Proportion Correct; PC) as well as with a reduction in performance accuracy (Learning Progress; LP).

In addition, we acquired objective measures of engagement: the pupillary response during task anticipation epoch of trials in the effort exposure phase. The pupil response has often been argued to reflect activity of the locus coeruleus (LC) (Murphy et al., 2014; Gilzenrat et al., 2010), the origin of noradrenaline, that is the neuromodulator most commonly implicated in task engagement (Aston-Jones & Cohen, 2005). Generally, an intermediate level of LC activity is considered optimal for task engagement, with both insufficient (boredom) and excessive arousal (stress) leading to impaired performance and reduced task engagement (Yerkes-Dodson curve (Yerkes & Dodson, 1908)). In fact, LC activity, and with it, the pupil response, has been shown to exhibit two modes of function: phasic and tonic. Phasic firing typically occurs in response to task-relevant events during epochs of high performance, and it is commonly characterized by large task-evoked dilations against a background of small baseline pupil size. This phasic mode is often contrasted with a tonic mode of pupil activity, which is associated with elevated baseline firing rate, absence of phasic responses, and degraded task performance (Aston-Jones et al., 1994; Aston-Jones & Cohen, 2005). By analogy, baseline pupil size was shown to be the highest and task-evoked dilations the lowest when participants decided to disengage from an effortful task (Gilzenrat et al., 2010) and this pattern reversed when participants reengaged with the task. Critically, in this prior work, decisions to engage were also always accompanied by higher accuracy, because participants were instructed to maximize accuracy-dependent reward. Therefore, participants chose to disengage from difficult tasks when their accuracy decreased too much.

Unlike this previous work on pupil dilation, the current task is predicted to disentangle effects of task accuracy and reported task engagement on pupil, with participants putatively reporting greater subjective task engagement for tasks on which they perform more poorly compared with tasks on which they perform well. Therefore, our paradigm provides a novel tool not only for testing the Learning Progress Motivation hypothesis of motivation (Oudeyer, Kaplan & Lopes, 2007), but also for investigating more directly the putative selective link between pupil dilation and task engagement. Because our task is designed to disentangle the linear pattern of task accuracy from the putatively nonlinear pattern of task engagement, the present study is also anticipated to substantiate prior assertions that pupil size reflects task engagement rather than task performance (Beatty, 1982; see van der Wel & van Steenbergen, 2018 for a review).

Thus, two straightforward, dissociable predictions are tested: First, during the cue period, the phasic pupil mode will track participants’ reported engagement, thus exhibiting a quadratic trend across task difficulty. Second, on a trial-by-trial basis, phasic pupil size will track proportion correct (PC) as well as learning progress (LP).

## 2. Methods

### 2.1. Participants

Forty English-speaking participants were recruited from the Radboud University participant pool (SONA Systems). All had normal or corrected-to-normal vision. All participants provided written informed consent to take part in the study, which was approved by the regional research ethics committee (Commissie Mensgebonden Onderzoek, Arnhem/Nijmegen, CMO2014/288). All participants received a monetary reward (€10.00) for their participation. Each participant was tested individually in a laboratory session lasting approximately 75 minutes. Participants were removed from the analyses if they had not completed the study (N = 2), or if no reliable pupil calibration accuracy could be obtained during the calibration phases (N = 2). The final sample size of 36 (ages 19-64; M = 24; *SD* = 9.6; 20 women see Supplementary Methods 1 for the age distribution and Supplementary Result 3 for analyses of a more homogeneous age-group) allowed us to detect a within-participant effect size of Cohen’s d ≥ 0.05 with 80% power and alpha of 0.05 (Cohen’s 1992).

### 2.2. Stimuli and Data Acquisition

During the computerized task portion of the experiment, participants were seated in a height-adjustable chair in front of a 23-inch monitor set to a resolution of 1920-1080 pixels, in a constant dimly lit room. Participants were instructed to keep their heads still and stabilized, rested in a chinrest positioned 50 centimetres away from the screen. All stimuli were delivered and controlled via the software Matlab (version 2016b) using Psychtoolbox library. Pupil responses were recorded using Eyelink 1000 eye tracker. Stimuli consisted of arithmetic summations of diverse difficulty levels, manipulated by summation length. Experimental scripts can be found on the author’s personal Github page (https://github.com/zceydas/OptimalEffort_Pupillometry).

### 2.3. Procedure

#### 2.3.1. The task

The participants started with a computerized task, which consisted of solving arithmetic summations varying in difficulty (e.g., 17 + 2). There were two different phases: a Capacity Phase to determine the four Task Difficulty levels, and a Performance Phase where those levels were performed. Both will be explained in detail in the following sections. In both phases, participants had to solve summations by giving a free response within a time period of 18 seconds. They were instructed to answer as many trials correctly as possible, and to answer as soon as they knew the correct answer. Participation rewards did not depend on task performance. The answer had to be entered digit by digit using numeric keys on the keyboard. Mistakes could be corrected using the “backspace” button. Their input was immediately displayed on the screen. All summations were presented in a row, with accurate feedback (*correct, incorrect* or *too late*) provided immediately afterwards.

#### 2.3.2. Capacity Phase

The Capacity Phase served to determine the participants’ level of skill. Each difficulty level consisted of 5 summation trials. Starting at the easiest possible level, the difficulty level of the summations was continuously increased by one level. Specifically, a level-up adjustment occurred when the participant correctly answered at least one question of the same difficulty level. This procedure was continued until the participant scored 0% correct at all 5 summations of a certain difficulty level. When the capacity phase was finished, a sigmoid function was fitted to the resulting datapoints in order to estimate four Task Difficulty levels based on participant’s own accuracy. The ‘Easy’ condition was kept the same across all individuals and yielded the following formula: X + X. All participants scored 100% correctly on this task condition. The subsequent task levels yielded lower accuracy: At ‘Intermediate1’ condition accuracy was 75%, at ‘Intermediate2’ condition accuracy was 50%, at ‘Difficult’ condition accuracy was 25%. These individually determined task conditions were used as the four Task Difficulty levels in the subsequent performance phase.

#### 2.3.3. Performance Phase

In the Performance Phase, the participant completed two blocks per Task Difficulty level in a pseudo-randomized order (Figure 1). Specifically, each Task Difficulty block was randomized in the first half of the performance phase (e.g. “C-A-D-B”), and the same order was repeated for the second half (“C-A-D-B”). Each block consisted of eight trials of one summation, so that the entire performance phase comprised sixty four trials in total. Prior to each block, participants underwent a standard nine-point calibration procedure of gazing shortly at nine markers displayed on the screen sequentially, to ensure pupil capture calibration accuracy was kept similar across task blocks. At the beginning of each trial, a letter cue (A, P, Q, Y) was presented to indicate the upcoming task condition. These cue-task condition pairings were counterbalanced across participants in such way that each letter had a similar likelihood of being paired with a task condition across participants. This cue period lasted three seconds, to capture the entire response profile of the pupil. The cue was followed by a fixation cross of one second. Following the fixation cross, the participant solved a summation trial, the difficulty of which depended on the current Task Difficulty block. All cues, fixation crosses and summations were presented in red. Feedback consisted of a blue check mark for correct, an orangish cross for incorrect and an orange clock for too slow. All stimuli were presented against a green background. Stimuli and background colours were selected to keep constant the screen luminance (0.3*R + 0.59*G + 0.11*B = ~110 cd/m^2^) as well as the contrast of the stimuli against the background during the whole task, to rule out any influence of screen luminance on pupil responses. Lastly, a final fixation cross appeared on the screen. Depending on response time (RT) in the summation period, the duration of this last fixation cross was adjusted, so that the total trial duration was always eighteen seconds. After each block, the participants provided a subjective flow rating, see below.

**Figure 1.**
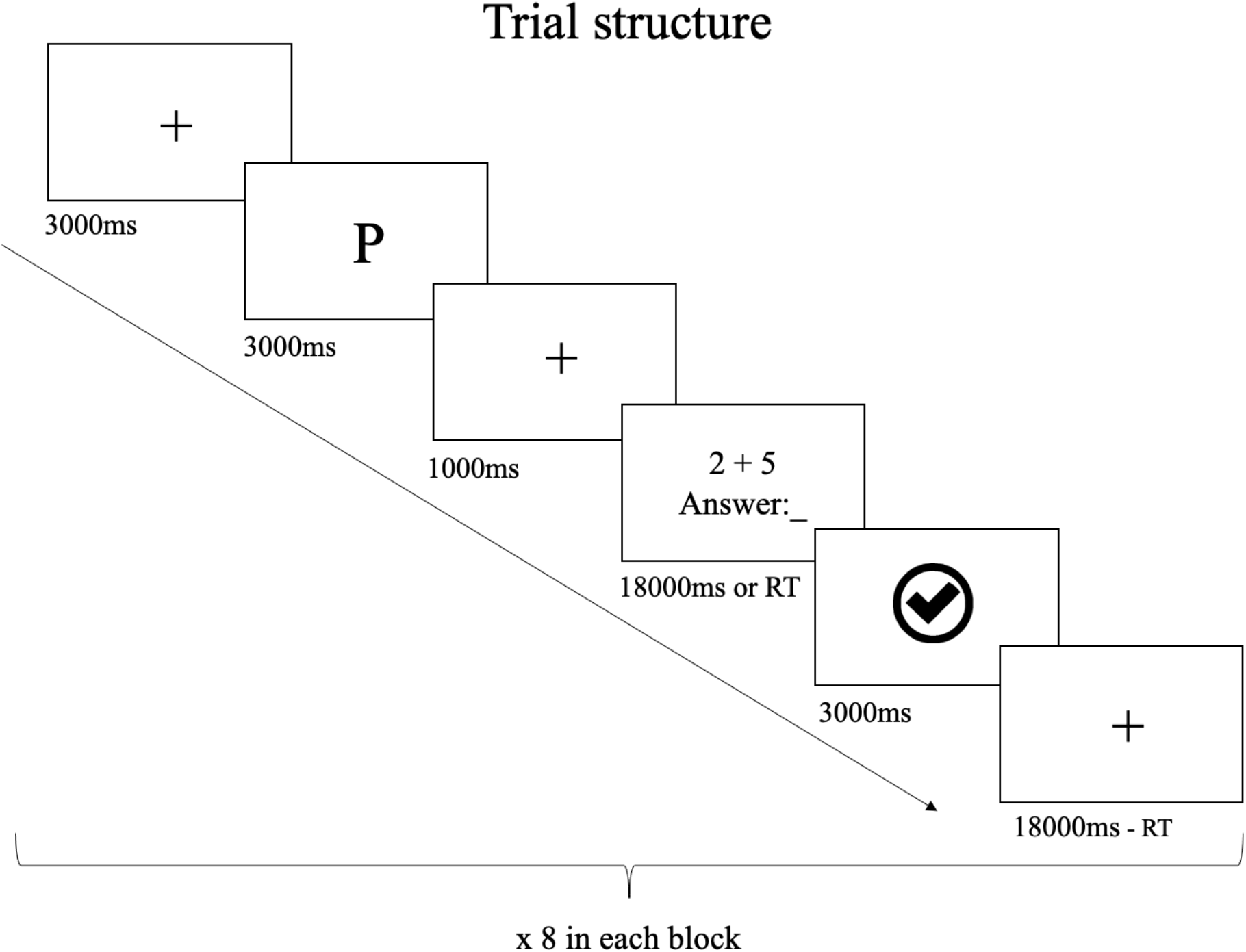
Illustration of one trial structure during the Performance Phase. The fixation cross preceding the cue presentation was 3 seconds. The cue was presented for 3 seconds and signalled the upcoming task performance in the Performance Phase. In the Capacity Phase, there was no cue. After the presentation of a fixation cross for 1 second, participants solved the arithmetic summation question. The deadline for this epoch was 18 seconds. Upon response, participants received accurate feedback for 3 seconds. The final fixation cross duration was variable depending on RT only in the Performance Phase. During the Capacity phase, the final fixation duration was randomized with a mean of 2 seconds.

#### 2.3.4. Subjective flow questionnaire

Subjective flow (flow index) was indexed by nine visual analogue ratings. The participant could range their response by moving the mouse on a horizontal line (10 cm in length) that had no anchors, except for the middle and endpoints. The answers were rated on a scale from 0 to 1. For 8 items, which measured the control component of flow, the endpoints were labelled *agree* and *disagree*. According to flow literature (Csikszentmihalyi, 1990; Ulrich et al., 2014; Ulrich, Keller & Grön, 2016), these items were used to monitor involvement, enjoyment, perceived fit between skills and task demands, and feeling of control with respect to each difficulty level (Table 1). A 9th statement assessed participants’ subjective sense of time (Keller & Bless, 2008; Ulrich et al, 2014).

**Table 1.**
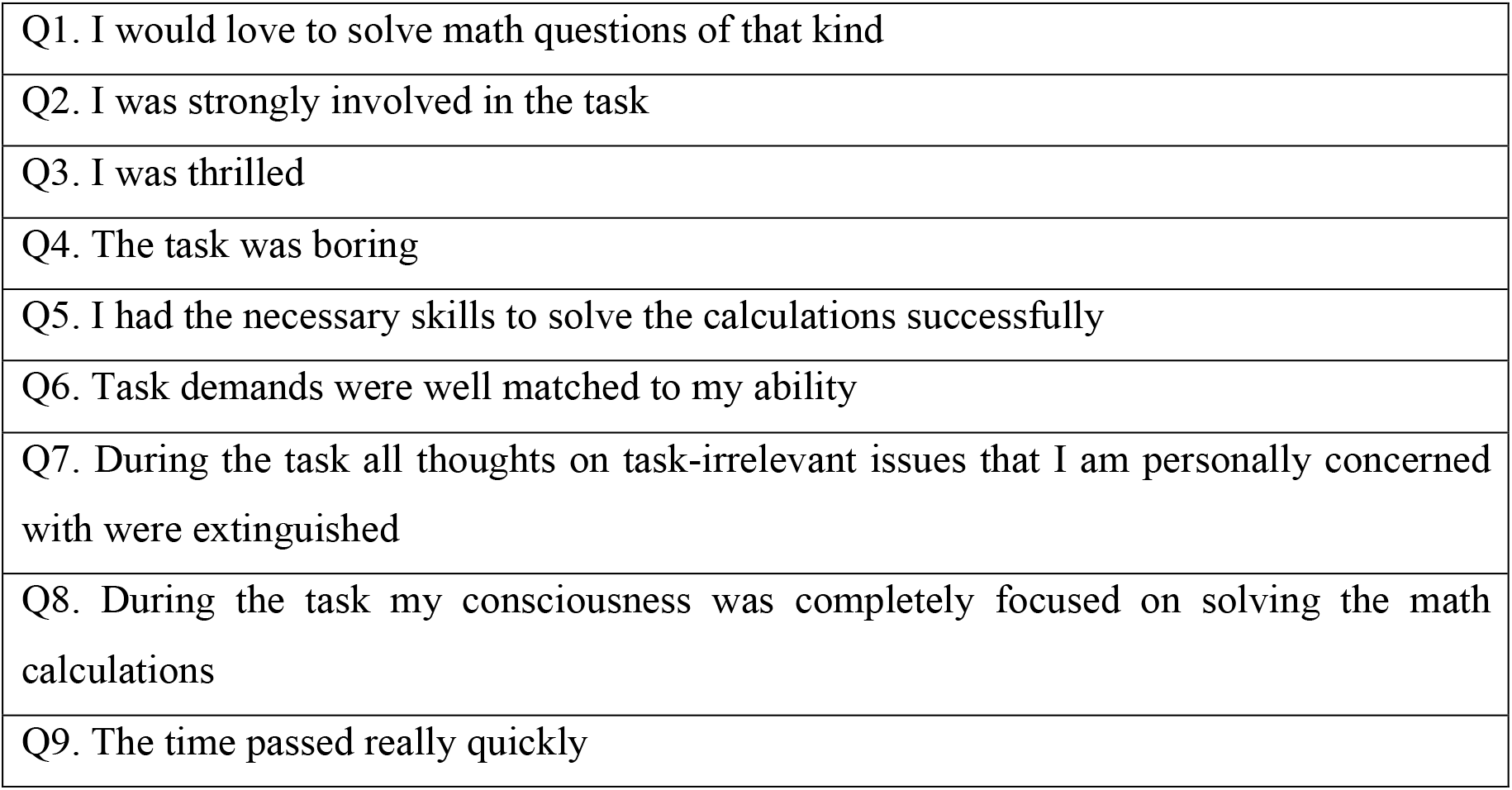
Flow questionnaire items.

### 2.4. Statistical analyses

All statistical analyses were conducted in Rstudio (Version 1.3.1093). The analyses included analysis of variance (ANOVA) and Bayesian modelling. For the ANOVA, we used the R package *ez*. Bayesian models were created in Stan and assessed with *brms* package (Bürkner, 2017). For ANOVA, alpha level of 0.05 was used for all analyses. For Bayesian models, credible intervals with 95% probability were computed and parameters that did not include 0 within their credible intervals were considered significant in predicting the outcome variable.

#### 2.4.1. Behavioural data analysis

##### 2.4.1.1. Response time and accuracy rate

To validate the paradigm, mean accuracy rates (percentage correct trials) and mean response times for correct trials (seconds) from the performance phase were submitted to a oneway repeated measures ANOVA using Task Difficulty level (easy, intermediate1, intermediate2, difficult) as a repeated measure. To assess simple effects of Task Difficulty, pairwise t-test analyses were executed with a Bonferroni correction. If the assumption of sphericity was not met, a Greenhouse-Geisser correction was applied (Field, 2009).

##### 2.4.1.2. Flow measurement analysis

To address whether participants’ flow ratings differ with Task Difficulty, we analysed the total flow score as well as four component scores (Supplementary Results 2): ‘ability’ (item 5 and 6), ‘involvement (item, 2, 7 and 8), ‘liking’ (item 1, 3 and 4) and ‘time’ (item 9). Individual ratings were averaged across all trials from each task condition, for total flow as well as for each of the four component scores. Total flow and component scores were submitted to a repeated measures ANOVA with Task Difficulty as a within-subject factor. Simple effects were investigated using Bonferroni corrected paired t-tests. In case of non-sphericity, a Greenhouse-Geisser correct was applied.

##### 2.4.1.3. Model parameter analysis

To understand the influence of trial-by-trial changes in accuracy and learning progress, we defined two latent parameters based on the recent work by Ten and colleagues (2021): proportion correct (PC) and learning progress (LP). PC was defined as a moving window average of accuracy over the last *n* trials of each block and updated across the two blocks for each Task Difficulty separately; LP was defined as the absolute value of the difference in PC over the previous and the current trial of that window. Therefore, PC is a moving average of accuracy over a time window (Figure 2) and LP reflects the absolute derivative of PC and indexes changes in task performance. Using the absolute value of LP implies that we consider both decrements as well as improvements in task performance. This definition of learning progress is based on the observation that lapses of attention and forgetting also motivate organisms to reallocate resources to tasks and aim to master tasks that are neither too easy nor too hard (Ten et al., 2021; Colas et al., 2019). In the case of our specific design, an effect of LP can thus reflect a boost of engagement on the trial after a lapse of attention.

**Figure 2.**
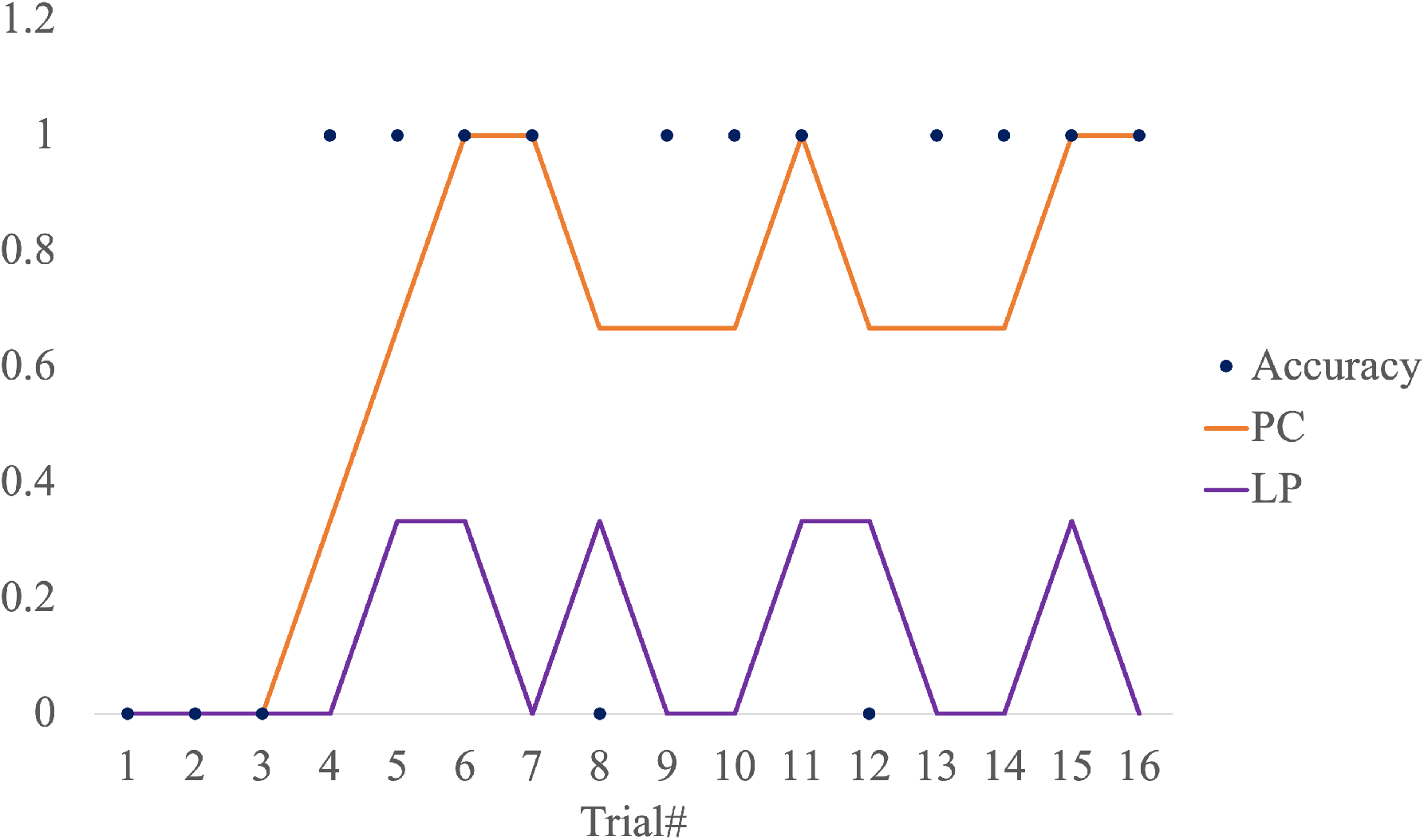
Exemplary progress of proportion correct (PC) and learning progress (LP) in relation to trial-by-trial changes in accuracy across all 16 trials (across two blocks) for the Intermediate1 level of one participant.

In order to determine the moving average window of PC, we considered three window sizes: 3, 4, or 5 trials. Consequently, we fit all PC and LP along with other variables of interest (*Task Difficulty, Session#, Trial#*, and *Task Order*) to predict pupil size in three separate models and compared their Bayesian Information Criterion (BIC) with the following formula notation: BIC = −2 * loglikelihood + d * log(N), where N is the sample size of the training set and d is the total number of parameters. Lower BIC scores is an indicator that the model is better. This revealed that the model that included PC over 3 trials explained pupil size better (BIC score of −3973.33) than did models that included 4 (BIC score of −3665.62) or 5 trials (BIC score of −3354.481). In order to demonstrate the functional form of these variables, we ran a repeated measures ANOVA with Task Difficulty as a within-subject factor. Significant effects were followed by pairwise Bonferroni corrected t-tests. In case of violation of the sphericity assumption, a Greenhouse-Geisser correction was applied (Field, 2009).

In order to ask whether if LP mediated the effect of task difficulty on flow scores, we first tested the effect of Task Difficulty on flow scores (X → Y), then tested the effect of Task Difficulty and LP on flow scores (X + M → Y) (Baron & Kenny, 1986) using *brms* function. In order to ask whether if LP mediated the effect of task difficulty on pupil size (see section 2.4.2 for pupillary data analysis), we first tested the effect of Task Difficulty on pupil size (X → Y), then tested the effect of Task Difficulty and LP on pupil size (X + M → Y) using *brms* models described above. If a mediation exists (full or partial), the effect of X on Y should diminish in the presence of M. Finally, the significance of the mediation analysis is tested using the *mediation* function in *bayestestR* package in R.

#### 2.4.2. Pupillary data analysis

As in previous literature, pupil size, reported as pupil diameter, was registered during fixation, cue and feedback periods with a sampling rate of 500Hz. The obtained raw pupillometry data were exported and pre-processed in Matlab before calculating trial-by-trial cue-related pupillary response during letter cue presentation. First, across all task epochs (1^st^ fixation, cue presentation, 2^nd^ fixation and performance feedback epochs) we excluded 12.1% of the trials from further analysis based on our exclusion criterion of more than 40% NaNs per trial (*M* = 7.72%, *SD* = 5.96%). These reflected blinking. In the remainder of the data, missing data and eye blinks were detected (*M* = 5.8% of overall data, *SD* = 4.48%), removed and smoothed by convolution with a 11 ms hanning-window. The smoothed pupil recordings were corrected using cubic spline interpolation. After interpolation, the pupil time series were low pass filtered using a 10 Hz first-order Butterworth filter to decrease noise (Knapen et al. 2016). Next, we checked for effects of Task Difficulty on gaze drift in x- and y-direction with a linear mixed effects model. This gaze drift control analysis confirmed that the frequency of the saccades in both directions did not change as a function of Task Difficulty (GazeDrift_X_: (*F*(3,34.33) = 0.89, *p* = .35); GazeDrift_Y_: (*F*(1,34.32) = 0.99, *p*=.32)). Therefore, the effect of saccades from pupil responses were not removed. As task difficulty was the same within the same task block, we did not use a baseline correction (Eckstein, Starr & Bunge, 2019; Hershman & Henik, 2019) on a trial-by-trial basis. As the cue presentation signalled the upcoming effortful task performance, the pupil response was calculated as the mean pupil diameter observed between a period of 2000 ms and 3000 ms after cue presentation (the end of the cue period). We chose this time window as previous research suggested that later time windows might reflect task preparation as pupil responses emerge slowly (Irons et al., 2017) and Figure 6 shows that this time window captures peak pupil dilation for all difficulty levels (Rondeel et al., 2015). Additional definitions for pupil size did not qualitatively change the results and are presented in Supplementary results.

The pre-processed data were transported to Rstudio. The trial-by-trial pupil size were rescaled between 0 and 1 and analysed using a Bayesian mixed effects regression model (brms on R). The model predictors included the following covariates of interest: the latent model parameters proportion correct (PC), learning progress (LP); as well as covariates of no-interest: Task Difficulty level, which refers to the difficulty of the task block ranging from 1 through 4, trial number and task order, where trial number refers to the trial number within a task block (1 through 8) and task order refers to the order in which a task block is presented to the participant among other blocks in the experiment (1 through 8).

#### 2.4.3. Analysis of pupil – behaviour links

All parameters were estimated using Monte Carlo (MHC) with 5 chains of 2000 samples each. Significant predictors are determined by looking at the posterior distribution; the evidence for an effect was considered significant if the posterior distribution contained zero within its range. All analyses were repeated using pupil size prior to cue onset (baseline) (pupil diameter during the 200 ms period before cue onset, instead of pupil response to the cue epoch). These revealed qualitatively the same associations between pupil and trial-by-trial changes in task performance (see Supplementary Results 4), indicating that anticipatory task-related pupil size was sustained throughout the blocks of same task difficulty. Task order, session number and trial number were included as nuisance variables in all relevant mixed effects models.

##### 2.4.3.1. Effect of task related factors on pupil size

To probe the effect of task-related factors on pupil size, we separately examined the influence of PC, LP on pupil size on a trial-by-trial basis. We used a Bayesian model comparison analysis of the non-averaged data, where subject-level parameters are drawn from group-level distributions. The model included pupil size as dependent variable, with fixed effects and random slopes for PC, LP, as well as nuisance variables Task Difficulty, Trial Number, Session Number and Task Order and Subject Number as random effect (model formula notation = (*PupilSize* ~ PC + LP + TaskDifficulty + trialNo + SessionNo + TaskOrder + (1 + PC + LP + TaskDifficulty + trialNo + SessionNo + TaskOrder || SubjectNumber, data=d, REML=F)).

Prior to each model fitting, predictor multicollinearity was assessed by computing variance inflation factors (VIF), which measures the inflation of a regression coefficient due to collinearity between predictors (Bruce & Bruce, 2017; James et al., 2014). For example, a VIF above 5 is considered problematic and predictors that yield problematic VIF scores are considered redundant. For this model, variance inflation factor (VIF) index for all predictors was low (VIF_TaskDifficulty_ = 1.09, VIF_PC_ = 1.08, VIF_LP_ = 1.03, VIF_TaskOrder_ = 1.04, VIF_SessionNumber_ = 2.00, VIF_TrialNo_ = 2.00).

##### 2.4.3.2. Motivational relevance of pupil size

To test the motivational relevance of pupil size, we asked whether flow scores can be predicted by pupil size. To this end, we adopted a Bayesian mixed effects regression model approach to assess the non-averaged data. Note that flow questionnaire responses were collected at the end of each task block and not on a trial-by-trial basis. Accordingly, the following model did not include the nuisance regressor *trial number*. The models included total flow index as the dependent variable, with fixed effects as well as random slopes for pupil size, and model parameters, PC and LP as well as nuisance variables Task Difficulty, task order, where Subject Number was entered as random effect (formula notation = (Flow ~ PupilSize + PC + LP + TaskDifficulty + SessionNo + TaskOrder + (1 + PupilSize + PC + LP + TaskDifficulty + SessionNo + TaskOrder || SubjectNumber, data=d, REML=F)). Variance inflation factor (VIF) index for all predictors was low (VIF_PupilSize_ = 1.01, VIF_TaskDifficulty_ = 1.01, VIF_PC_ = 1.01, VIF_LP_ = 1.00, VIF_TaskOrder_ = 1.01, VIF_SessionNumber_ = 1.01, VIF_TrialNo_ = 1.01), justifying the inclusion of these terms in the same model.

Model fitting was followed by mediation analyses. We asked whether if LP mediated the effect of pupil size on flow scores, we first tested the effect of pupil size on flow scores (X → Y), then tested the effect of pupil size and LP on flow scores (X + M → Y). Finally, the significance of the mediation analysis is tested using the *mediation* function in *bayestestR* package in R.

##### 2.4.3.3. Behavioural relevance of pupil size

To test the behavioural relevance of pupil size, we assessed whether task performance can be predicted from pupil size, again using a Bayesian mixed effects regression model of the non-averaged data. The two separate models included current trial accuracy and RT as dependent variables, with fixed effects as well as random slopes for pupil size, and model parameters, PC and LP and nuisance variables Task Difficulty, trial number and task order, where subject number was entered as random effect (formula notation = (*TaskPerformance* ~ PupilSize + PC + LP + TaskDifficulty + SessionNo + TaskOrder + (1 + PupilSize + PC + LP + TaskDifficulty + SessionNo + TaskOrder || SubjectNumber, data=d, REML=F)).

For the model predicting accuracy, the variance inflation factor (VIF) index for all predictors was low (VIF_PupilSize_ = 1.03, VIF_TaskDifficulty_ = 1.46, VIF_PC_ = 1.41, VIF_LP_ = 1.07, VIF_TaskOrder_ = 3.98, VIF_SessionNumber_ = 4.19, VIF_TrialNo_ = 1.23), justifying the inclusion of these terms in the same model. For the model predicting correct RT, the variance inflation factor (VIF) indices for all predictors were low (VIF_PupilSize_ = 1.03, VIF_TaskDifficulty_ = 1.46, VIF_PC_ = 1.46, VIF_LP_ = 1.08, VIF_TaskOrder_ = 3.62, VIF_SessionNumber_ = 3.79, VIF_TrialNo_ = 1.21), permitting the inclusion of all predictors in predicting correct RTs.

## 3. Results

### 3.1. Effect of Task Difficulty on task performance

Confirming our experimental manipulation, accuracy rate significantly decreased with Task Difficulty (*F*(3,105) = 106.639, *p* < .001) (Figure 3A). Accuracy at each Task Difficulty level was significantly different from the others (all *p*s<0.001).

**Figure 3.**
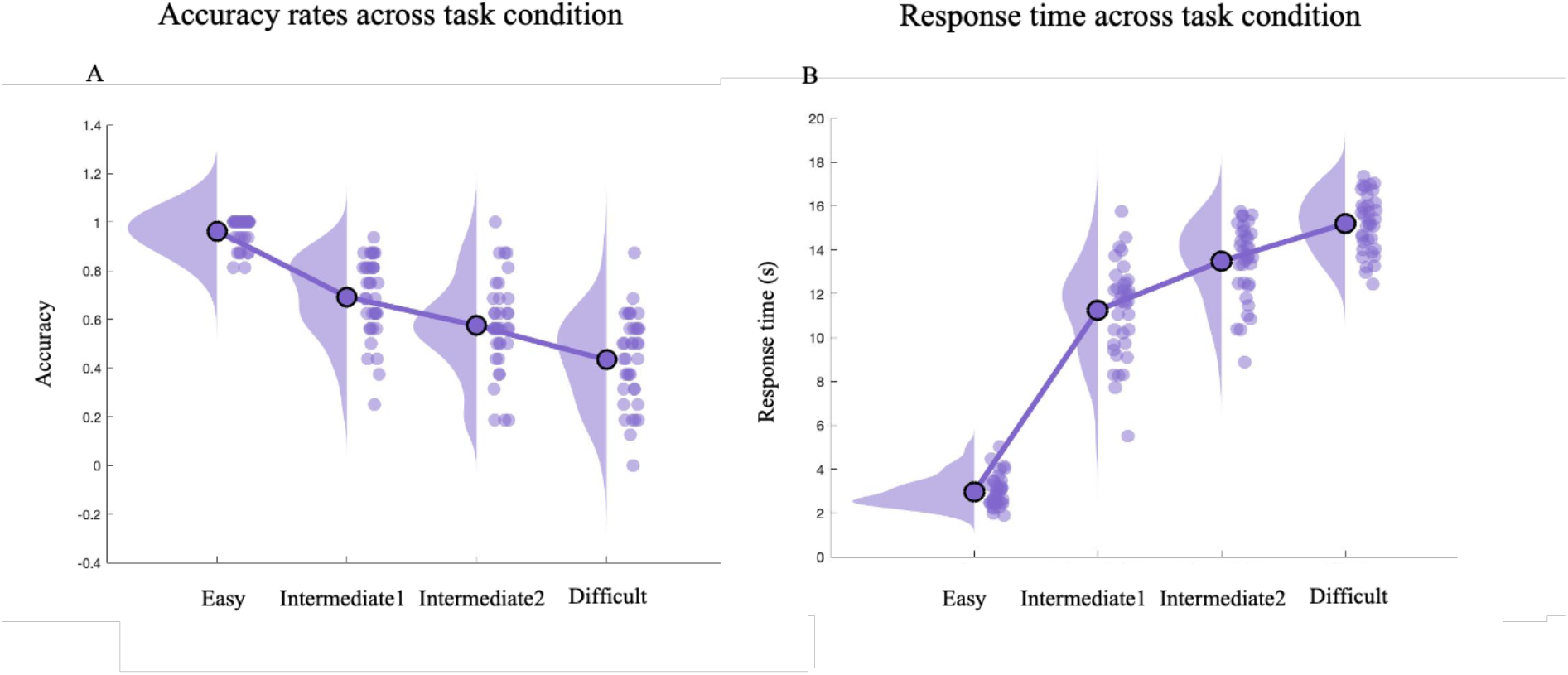
Raincloud plot for A) accuracy rate, B) response time (RT) across task condition. The dots display each participant’s mean accuracy rate and correct RT. Circled dots stand for group averages.

Similarly, the time it took to respond correctly increased with Task Difficulty (*F*(2.267,79.335) = 755.848, *p* < .001) (Figure 3B). Average RT at each Task Difficulty level was significantly different from the others (all *p*s<0.001).

### 3.2. Effect of Task Difficulty on subjective flow scores

Although task performance declined monotonically with Task Difficulty, subjective engagement as indexed by the flow questionnaire increased with Task Difficulty. The flow index significantly differed across Task Difficulty (*F*(1.686,140) = 19.186, *p* < .001) (Figure 4). Both intermediate difficulty levels yielded greater flow scores compared with the Easy level (both *p*s<0.001) and Difficult yielded greater flow scores compared with the Easy level (p<0.001). No significant effects differences were found between Intermediate1 and Intermediate2 (*p* = 1.000), Intermediate1 and Difficult (*p* = 0.412, nor Intermediate2 and Difficult (*p* = 1.000), suggesting that the subjective experience of flow plateaued at the intermediate Task Difficulty levels. A closer look into the subcomponents of the flow inventory showed that task involvement and liking increased with Task Difficulty (see Supplementary Results 1), while perceived task ability decreased, indicating that task engagement, as indexed by the subjective experience of flow, dissociated from that of task ability.

**Figure 4.**
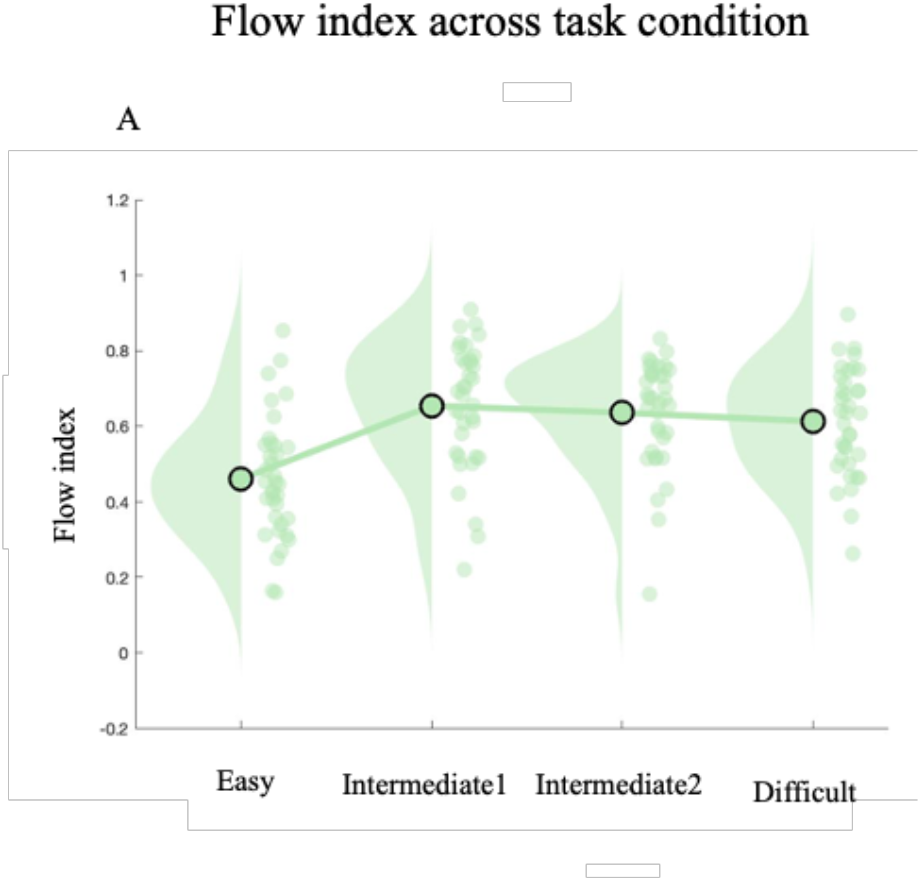
Raincloud plot for average flow scores across Task Difficulty. Circled dots stand for group averages.

### 3.3. Effects of Task Difficulty on latent model parameters (proportion correct (PC) and learning progress (LP)), pupil size and flow scores

Average proportion correct (PC) (Figure 5) declined significantly with increasing Task Difficulty (*F*(3,36.57) = 94.76, *p* < 0.001) where the pairwise difference in PC between all tasks were significant (all *p*s<0.001).

**Figure 5.**
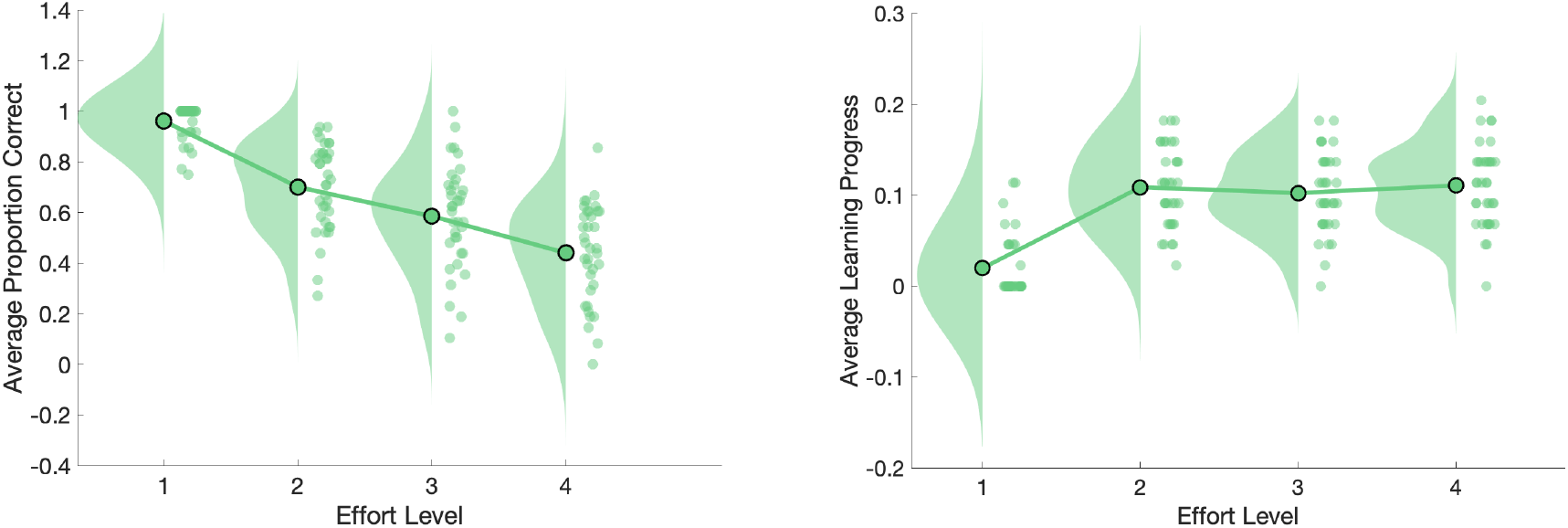
Raincloud plot for A) Average PC, and B) Average LP. The dots display each participant’s mean PC and LP. Circled dots stand for group averages.

Learning progress (LP) magnitude differed significantly as a function of Task Difficulty (*F*(3,55.24) = 36.95, *p* < 0.001), but plateaued at the intermediate difficulty levels (Figure 5). Both Intermediate Task Difficulty levels and the Difficult level yielded greater LP compared with Easy (all *p*s <0.001), but there was no difference between intermediate levels and Difficult or Intermediate1 vs Intermediate2 (Intermediate1 vs Intermediate2, *p* = 0.16); Intermediate1 vs Difficult, *p* = 0.37; Intermediate2 vs Difficult, *p* = 0.99).

To assess whether LP mediates the effect of Task Difficulty on flow scores, we ran mediation analysis. The total effect of Task Difficulty on flow scores (B = 0.409, CI [0.184, 0.635]) was significantly smaller in the presence of LP (0.397, CI [0.173, 0.624]), indicating that the effect of Task Difficulty on flow scores was partially mediated by LP (B = 0.013, CI [0.000, 0.026]).

To assess whether LP mediates the effect of Task Difficulty on pupil size, we ran mediation analysis. The total effect of Task Difficulty on pupil size (B = 0.018, CI [0.011, 0.024]) was significantly smaller in the presence of LP (0.017, CI [0.010, 0.023]), indicating that the effect of Task Difficulty on pupil size was partially mediated by LP (B = 0.001, CI [0.000, 0.002].

### 3.4. Effects of learning progress on pupil size

To assess which (trial-by-trial latent) processes/factors contribute to trial-by-trial changes in pupil size, we ran a mixed effects model with predictors of interest, *PC, LP* and the nuisance regressors*, Task Difficulty, trial number, session number* and *task order*.

There were significant effects of PC (B = −0.03, CI [−0.05, −0.01]) and LP (B = 0.03, CI [0.01, 0.05])) on pupil size. Specifically, pupil response was greater in trials following lower average accuracy and greater Learning Progress (Figure 6, 7A).

**Figure 6.**
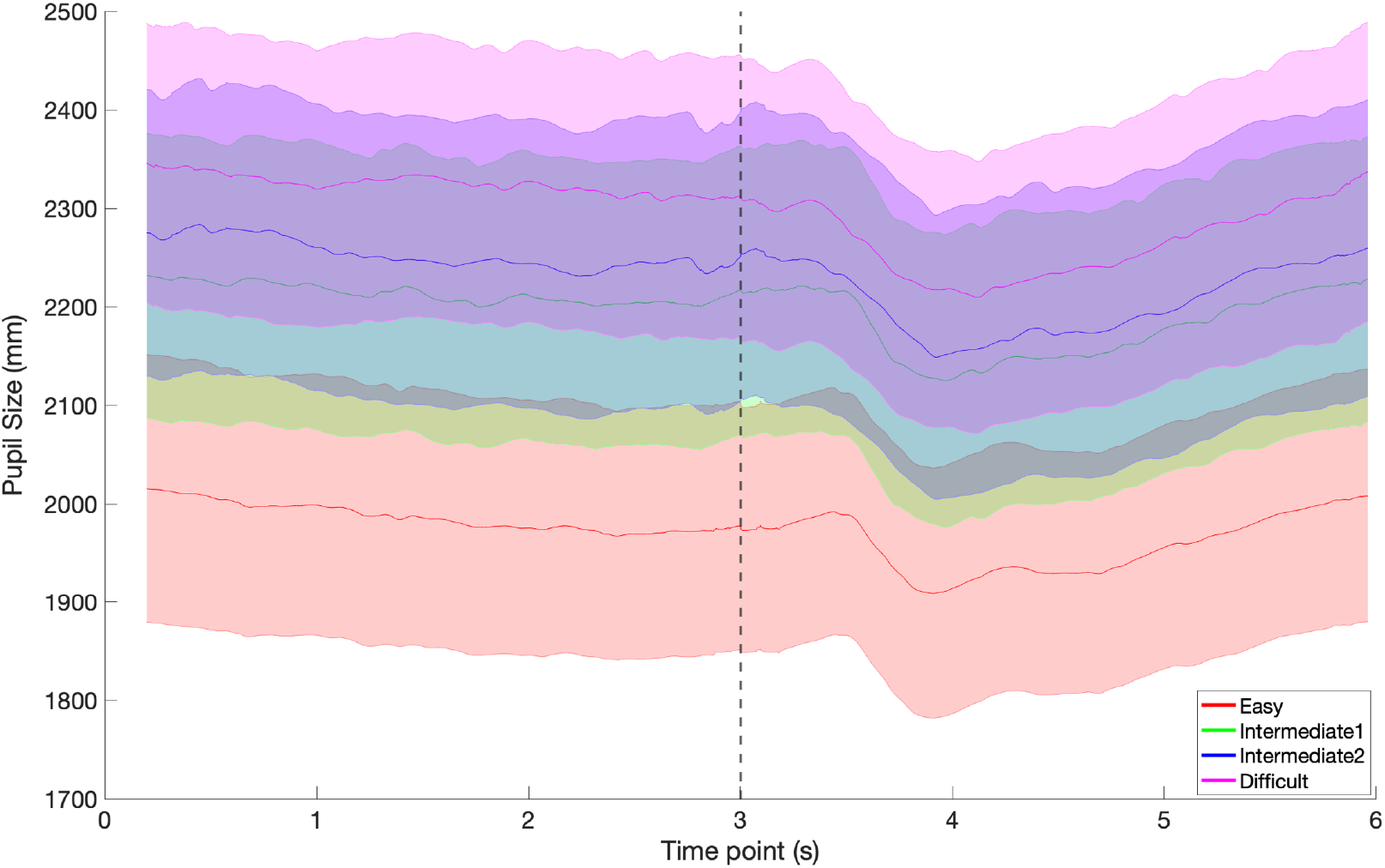
Average time course of the pupil size by Task Difficulty. Dashed line indicates cue onset.

**Figure 7.**
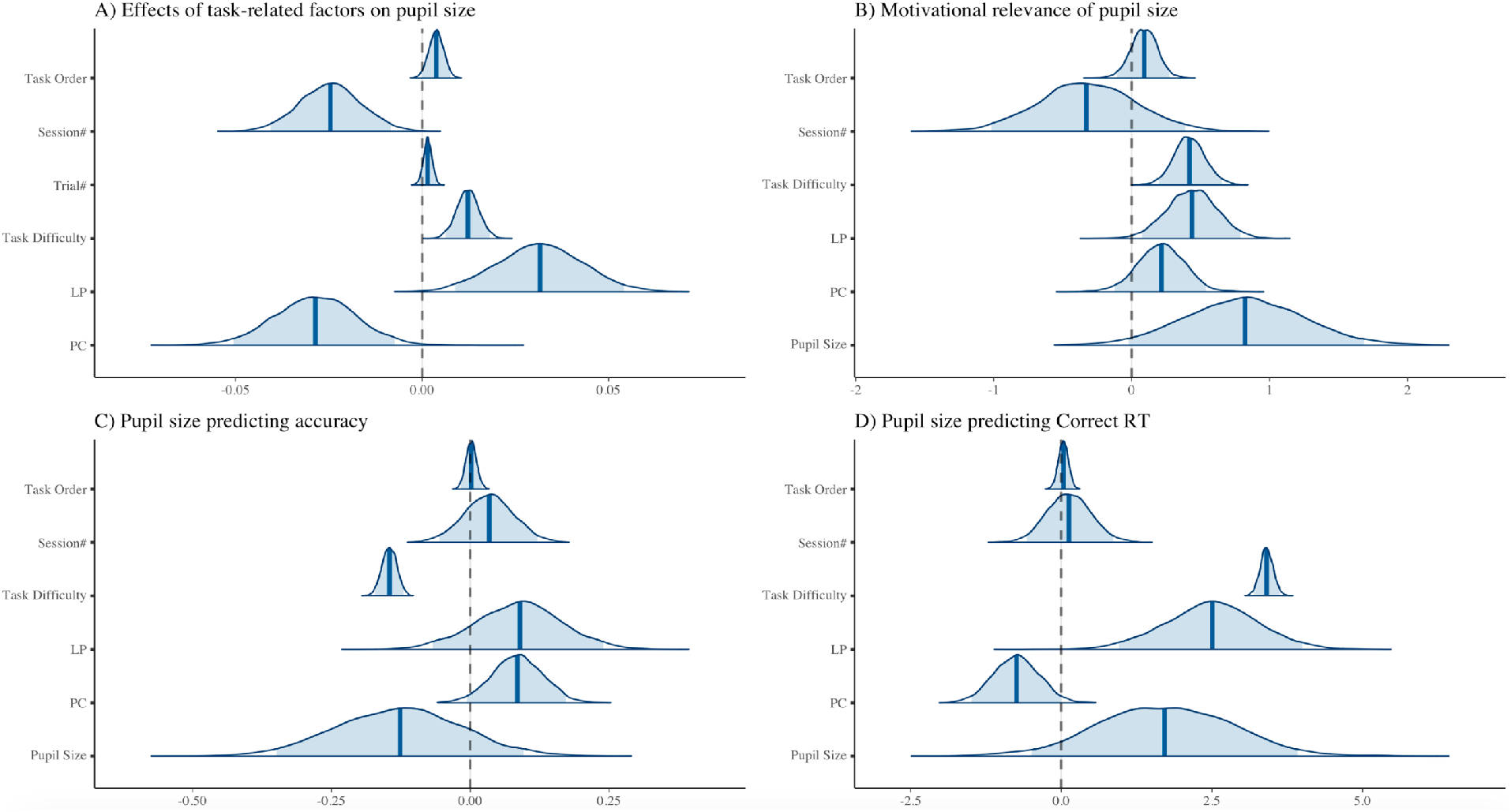
Densities of model parameter estimates. A) Model parameter densities for predicting pupil size. B) Model parameter densities for predicting flow scores. C) Model parameter densities for predicting accuracy. D) Model parameter densities for predicting Correct RTs.

### 3.5. Effect of pupil size and learning progress on subjective flow ratings

Next, we assessed the motivational relevance of pupil size by testing whether pupil size predicts subjective flow scores.

The model results (Figure 7B) showed participants reported greater flow scores with (marginally) greater pupil size (B = 0.82, CI [−0.02, 1.67]) and greater LP (B = 0.43, CI [0.06, 0.78]).

To assess whether LP mediates the effect of pupil size on flow scores, we ran mediation analysis. The total effect of pupil size on flow scores (B = 0.836, CI [−0.035, 1.694]) was smaller in the presence of LP (0.817, CI [−0.045, 1.671]), although this effect was not significant (B = 0.020, CI [−0.011, 0.066], indicating that subjective flow scores could be differentially accounted by pupil size and learning progress.

### 3.6. Effects of pupil size on task performance

Finally, we assessed the behavioural relevance of pupil size by testing the influence of pupil size on upcoming task accuracy and correct trial RTs. The full model included the predictors of interest, *pupil size, PC, LP*, and the nuisance regressor *Task Difficulty, task order* and *session number*.

The model results predicting accuracy (Figure 7C) showed that the predictors of trial-by-trial accuracy was PC (B = 0.09, CI [0.00, 0.17]), while the effect of LP and pupil size were insignificant (pupil size: B = −0.13, CI [−0.36, 0.09]; LP (B = 0.09, CI [−0.08, 0.24]). As expected, accuracy was higher following trials with greater PC.

The model results predicting correct RTs (Figure 7D) showed that the predictors of trial-by-trial RT were PC (B = −0.75, CI [−1.48, −0.01]) and LP (B = 2.50, CI [0.99, 4.03]) while the effect of pupil size was insignificant (B = 1.70, CI [−0.49, 3.93]). Thus, correct trial RTs were longer on trials following smaller PC and greater LP.

## Discussion

While a substantial body of work has shown that cognitive effort is aversive and costly (Kool et al., 2010; Westbrook, Kester & Braver, 2013; Sayali & Badre, 2020), a separate line of research on intrinsic motivation suggests that people spontaneously seek challenging but learnable tasks. According to one prominent account of intrinsic motivation, the Learning Progress Motivation theory, the preference for difficult tasks reflects the dynamic range that these tasks yield for changes in task performance (Ten et al., 2021; Kaplan & Oudeyer, 2007; Oudeyer, Gottlieb & Lopes, 2016). Here we tested this hypothesis, by asking whether greater engagement with cognitive effortful tasks, indexed by subjective ratings and objective pupil measurements, is a function of trial-wise changes in task performance.

Across four individually assigned difficulty levels, the current study dissociated subjective task engagement from task difficulty. As such, the current results showed that subjective engagement and liking scores, as indexed by the flow questionnaire, increased with increasing task difficulty despite significant decreases in task performance and perceived task ability (Supplemental results 2). By leveraging a design from previous studies on flow states (Ulrich et al., 2014; Ulrich, Keller & Grön, 2016; Katahira et al., 2018), we showed that difficult task levels yielded greater LPs compared with the easiest difficulty level, leading to a plateau of task performance increases as well as subjective engagement scores at the intermediately difficult task levels. This plateau effect stands in contrast to previous flow findings (Ulrich et al., 2014; Ulrich, Keller & Grön, 2016), which showed an inverted-U shape function of subjective task engagement across difficulty levels. We argue that this discrepancy might be due to differences in study designs. We found that learning progress increases with greater task difficulty (plateauing at Intermediate 1) because participants showed greater task improvement at the most difficult task level which they were initially around 25% correct, increasing average accuracy of that level by ~40% accuracy across two blocks. Thus, the most difficult task was still within the capacity range of the participants, and hence, was not impossible for them to master. This is in contrast with Ulrich et al. (2014, 2016)’s ‘overload’ task manipulation in which the task level was way above the participant’s own capacity and the average accuracy rate was around 5%. This difference between paradigms made the most difficult task condition in our study another intermediately challenging task condition. Importantly, we found that subjective engagement also increased with increasing effort level and learning progress, where the link between cognitive effort and subjective flow scores could be partially attributed to learning progress. Thus, we have shown that mentally more challenging tasks which yielded the greatest range for task performance changes received greater liking and engagement scores than the easiest level, leading to plateau of subjective pleasure across difficulty levels. These results suggest that what might be underlying the subjective pleasure during flow states might be the associated increased dynamic range for changes in task performance.

This observation is generally consistent with previous literature indicating that perceived effort costs can be alleviated by subjective reward received during task performance (Inzlicht, Shenhav & Olivola, 2018). Furthermore, Devine and Otto (2021) have shown that receiving temporal information about progress reduced demand avoidance in a demand selection task in the absence of reward, indicating that information regarding task performance influenced cost-benefit decisions regarding cognitive effort. These results underscore the value of performance progress in mediating effort costs. As such, performance variability, as tracked by learning progress variable in our study underlie subjective and objective measures of engagement. This prediction concurs with the finding that (Baranes, Oudeyer & Gottlieb, 2014), in a video game setting, when people are given the option to either repeat the same difficulty level or to voluntarily increase difficulty, they gradually increase the difficulty of the game they play even when it means having poorer task performance. Consistently, Ten and colleagues (2021) showed that participants who were motivated to maximize learning in task activities showed greater tolerance for their own performance errors and better learning outcomes. The current study corroborates these observations and firmly establishes the link between effortful task engagement and changes in task performance as a function of learning progress.

Our findings are reminiscent of studies suggesting a key role for efficacy in the willingness to exert cognitive effort. Efficacy refers to how much one’s invested efforts will impact the outcome of these efforts (Bandura, 1986). For example, if an outcome is driven mostly by factors outside of one’s control, one’s efficacy would be low. An updated version of the Expected Value of Control theory (EVC; Shenhav, Botvinick & Cohen, 2013) incorporates self-efficacy as a determinant of cost-benefit calculations regarding effort allocation, where higher efficacy increases the amount of effort exertion (Blain & Sharot, 2021; Frömer et al., 2021; Shenhav, Botvinick, & Cohen, 2013). Consistent with this framework, a recent study manipulated subjective efficacy by changing the contingency between actions and reward outcomes in a Stroop task design (Frömer et al., 2021). They demonstrated that if participants perceived their efforts as efficacious, they are more likely to exert control, as indexed by higher contingent negative variation (CNV) amplitude, an event-related potential (ERP) known to track proactive control allocation, and higher P3b, ERP known to signal incentive evaluation, during initial cue period. The current study goes beyond that prior work by substantiating Learning Progress Motivation theory, according to which changes in task performance registers as intrinsic value. These results might thus inspire the updating of current resource allocation models of cognitive effort, such as EVC, with parameters capturing changes in task performance.

The current study results raise the possibility that increases in challenging task engagement are accompanied by task performance-based increases in arousal, perhaps mediated by locus coeruleus (LC) activity and noradrenaline. According to Adaptive Gain Theory of LC (Aston-Jones & Cohen, 2005), there are two modes of LC activity: tonic and phasic. Baseline and shortterm burst-like activities of LC are typically inversely correlated. In a tonic mode, baseline LC activity is greater than momentary LC bursts. In this mode, the system is disengaged, and behavior is inattentive or distractible. On the other hand, the phasic mode is characterized by low baseline LC activity and greater LC bursts that are typically coupled with task-relevant outcomes or responses. Thus, in order to verify the subjective preference of challenging tasks, we related cue evoked pupil size, as an objective marker of task engagement to subjective engagement scores.

Previous research has shown that phasic pupil response, which has been demonstrated to correlate with LC activity (Gilzenrat et al., 2010), tracks task utility where reward was coupled to task performance. The results were interpreted to reflect disengagement from tasks that were difficult. In contradiction with these observations, the current results show that easier tasks where expected accuracy was higher, were accompanied by smaller pupil size in the absence of performance-contingent rewards. As such, previous research (Massar et al., 2016) showed that an increase in pupil diameter was only present when rewards were contingent on good performance (high reward condition) but not when reward was provided at random (random reward condition), indicating the coupling between pupil size and expected reward might be mediated by subjectively felt contingency of reward. The current result show that in the absence of performance-contingent rewards, pupil size increases with increasing task difficulty, implying the intrinsic utility of mental challenge.

More critically, pupil size was negatively predicted by proportion correct (PC) and positively predicted by learning progress (LP). Previous evidence indicates that pupil size increases monotonically with greater task difficulty (Belayachi et al., 2015; Brouwer, et al., 2014; Irons, Jeon & Leber 2017; Klingner, Tversky & Hanrahan, 2011; Moresi et al., 2008), although the link between task accuracy and pupil size has been attributed to task engagement rather than task difficulty alone (van der Wel & van Steenbergen, 2018). Our results consistently demonstrated that task-evoked pupil size increases with task difficulty as well as with variability in task accuracy (LP). As such, the effect of task difficulty on pupil size was partially mediated by learning progress, indicating that the link between task difficulty and pupillary measures of engagement can be attributed to the variability in average task accuracy. Moreover, pupil size is predictive of subjective scores of engagement, as indexed by flow scores in the current study, indicating that pupil size tracks task engagement. This conclusion is generally in line with recent evidence presented by da Silva Castanheira, LoParco, and Otto (2020), who demonstrated that task-evoked pupil size changes as a function of effort investment, even when task demands were kept constant. Their finding that pupil size was associated with better task performance, despite constant task demands, led them to propose that the pupil response serves as a reliable index of cognitive effort investment. Here we go beyond this by showing that this link exists even in the presence of accuracy decline, and can be accounted for, in part by changes in learning progress (changes in performance accuracy).

Importantly, the current study tested the relationship between task difficulty and pupil size during a cue period and not during task performance. As such, pupil size tracked the average accuracy of the upcoming task, consistent with previous accounts (Kurniawan, Grueschow & Ruff, 2021) which dissociated the time of effort anticipation from that of effort exertion in an instrumental effort paradigm and showed that pupil size tracked the anticipation of the upcoming effort level of the task in a voluntary choice paradigm. Moreover, these increases in pupil dilation were stronger when the participants accepted to exert the effort option versus not, suggesting that pupil size might be associated with the energization required to perform an upcoming action. The current results, combined with the previous literature, suggests that cue evoked pupil size signals expectations about effort requirements rather than the exertion of effort itself.

The current results corroborate these findings by showing that pupil size tracked performance changes independent of task difficulty, potentially boosting task engagement in more challenging but learnable tasks via noradrenergic arousal mechanisms. The trial-by-trial relationship between performance change and pupil size, provides direct evidence for the Learning Progress Motivation theory (Oudeyer, Gottlieb & Lopes, 2016), which states that motivation for challenging tasks is a function of the opportunity for improving performance and learning.

These findings also parallel evidence that the experience of flow is characterized by an intermediate level of arousal, as indexed by intermediate sympathovagal system activation (heart rate and breathing rate) and an intermediate level of ACC activity (de Sampario Barros et al., 2018; Ulrich et al., 2014; Ulrich, Keller & Grön, 2016). However, no studies to date have directly tested the relationship between the LC-associated pupil response, performance changes and optimally challenging effort. The current study firmly established such a relationship between pupil size and challenging task anticipation by showing that pupil size predicted greater subjective task engagement as assessed by the flow questionnaire at the end of each task block as assessed by flow scores.

While the mechanism underlying intermediate effort preference in our study points to the role of LC-based arousal, pupil size has also been associated with dopaminergic activity (Manohar & Husain, 2015; Muhammed et al., 2016). The dopaminergic system is mostly located in the midbrain ventral tegmental area (VTA) and substantia nigra (SN), and is traditionally involved in reward processing and value-based behaviour (Olds & Milner, 2020; Schultz, Dayan & Montague, 1997; Schultz, 2007). Moreover, DA is involved in tracking reward uncertainty, in a way to facilitate learning (Gershman & Uchida, 2019; O’Doherty et al., 2003). Future studies should disentangle the role of noradrenergic and dopaminergic systems in mediating the relationship between intermediate challenge and pupil size.

The interpretation of the results also are not without constraints. As described earlier, unlike in the flow induction paradigm (Ulrich et al., 2014; Ulrich, Keller & Grön, 2016), our design did not include a supra-threshold capacity difficulty level. Thus, in our task, all task conditions except the easiest difficulty level could be mastered by the participant. Therefore, the current design cannot answer what factors underlie the (dis)engagement during tasks that are above capacity limits. Secondly, although we measured subjective and pupillary engagement in the absence of external reward, in order to induce a state of flow, we provided performance feedback (Csikszentmihalyi, 1990). Hence, we were not able to test whether these findings would also hold in the absence of external feedback, which might be considered a form of external reward, underlie individual variability in feedback sensitivity and complicate our definition of internal rewards.

## Supplementary Material

### Supplementary Methods 1

Distribution of participants’ ages was skewed (Supplementary Figure 1). The Shapiro-Wilk statistic associated with participants’ ages was W = 0.60, indicating that age distribution significantly deviated from normality (*p* < 0.001).

**Supplementary Fig 1.**
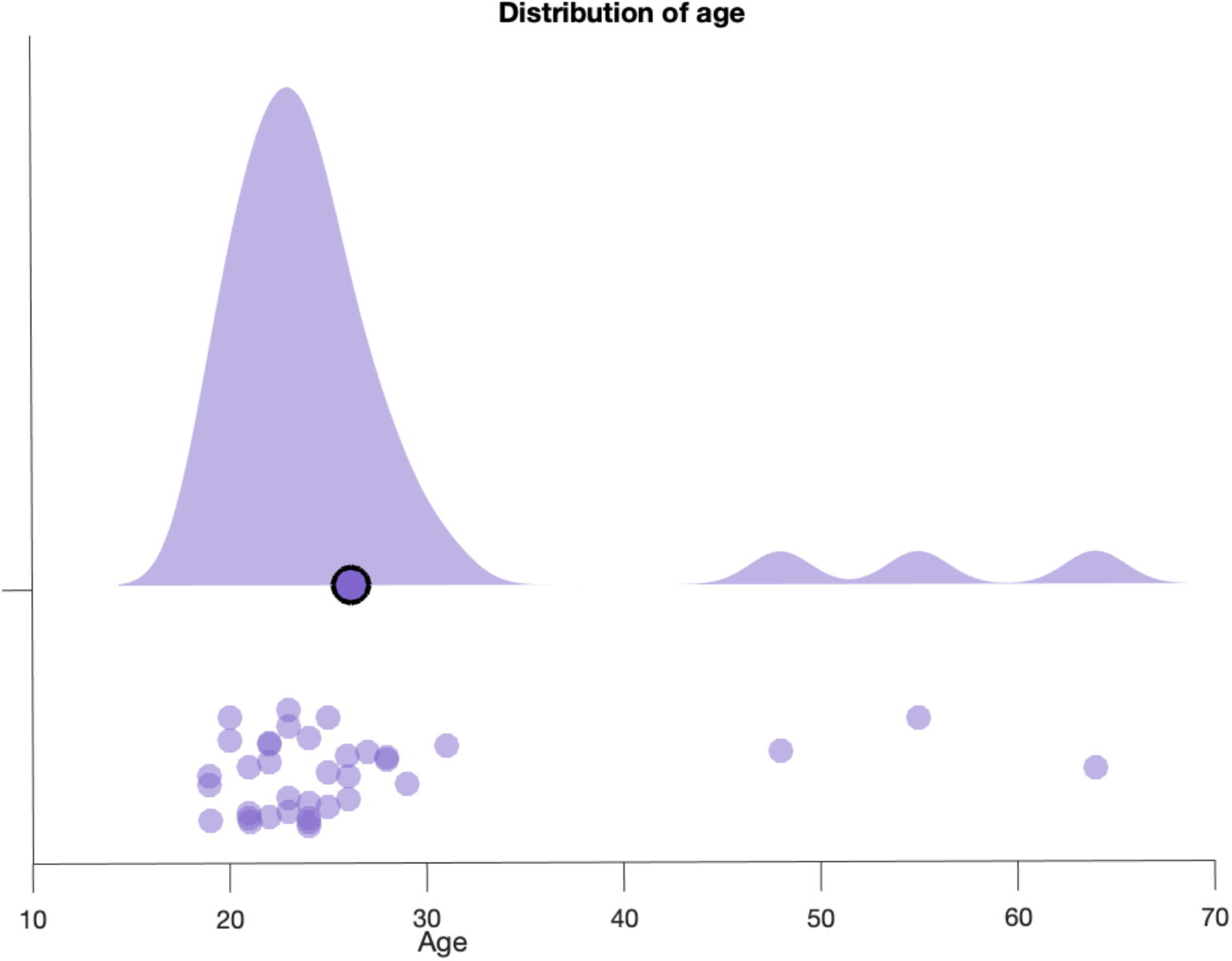
Distribution of participants’ ages. Circled dot stand for the group average.

### Supplementary Results 1

#### The effect of Task Difficulty on the subcomponents of Flow inventory

As the flow questionnaire consisted of components of perceived engagement, liking, ability and flow of time, we examined effects of Task Difficulty on each component separately (Supplementary Figure 2).

The results showed a significant effect of Task Difficulty on involvement (*F*(1.479,51,765) = 32.617, *p* < .001), where involvement scores significantly increased with Task Difficulty. Again, subjective engagement scores plateaued at intermediate Task Difficulty. Pairwise t-tests revealed significantly lower scores for easy versus Intermediate1 (*p* < .001), easy versus Intermediate2 (*p* < .001), easy versus Difficult (M = 0.720, SD = 0.035, *p* < .001). No significant differences were found for Intermediate1 versus Intermediate2 (*p* = 0.391), Intermediate1 versus Difficult (*p* = 0.120), nor Intermediate2 versus Difficult (*p* = 1.000).

Liking scores also significantly differed across Task Difficulty (*F*(2.379,83.265) = 11.148, *p* < .001), but this time the effect of Task Difficulty plateaued at Intermediate1 . Specifically, liking scores were significantly lower for Easy versus both Intermediate levels (Intermediate1: *p* < .001; Intermediate2: *p* = .005) and did not differ between Easy vs Difficult (*p* = 0.091), or Intermediate vs Difficult (between intermediate1 and intermediate2 (*p* = 0.339), Intermediate1 and Difficult (*p* = 0.114) nor between Intermediate2 and Difficult (*p* = 0.928)).

Consistent with their own task performance, participants perceived ability at each Task Difficulty level showed a declining trend. Subjective ability scores significantly different across Task Difficulty levels (*F*(2.313,80.955) = 69.095, *p* < .001). Specifically, Easy and Intermediate1 level yielded higher ability compared with Difficult Task Difficulty (Easy: p<0.001; Intermediate1: *p* = 0.005), while other Task Difficulty, including Easy and intermediate levels did not significantly differ from each other (easy vs. intermediate1 (*p* =1.000), easy vs. intermediate2 (*p* = 0.209)), indicating that participants rated their own ability for the easiest and intermediate difficulty levels similarly.

Lastly, perceived time on task showed a significant inclining linear trend across Task Difficulty (*F*(1.428,49.98) = 32.492, *p* < .001). Specifically, both intermediate difficulty levels yielded higher scores compared with Easy (both *p*s < .001), and easy compared with difficult (*p* < .001), while intermediate levels did not significantly differ from each other (*p* = 0.872) nor from Difficult (Intermediate1 vs. Difficult, *p* = 0.643; Intermediate2 vs. Difficult, *p* = 1.000), indicating that subjective time passed quicker with increasing difficulty levels and plateaued at intermediate difficulty.

**Supplementary Fig 2.**
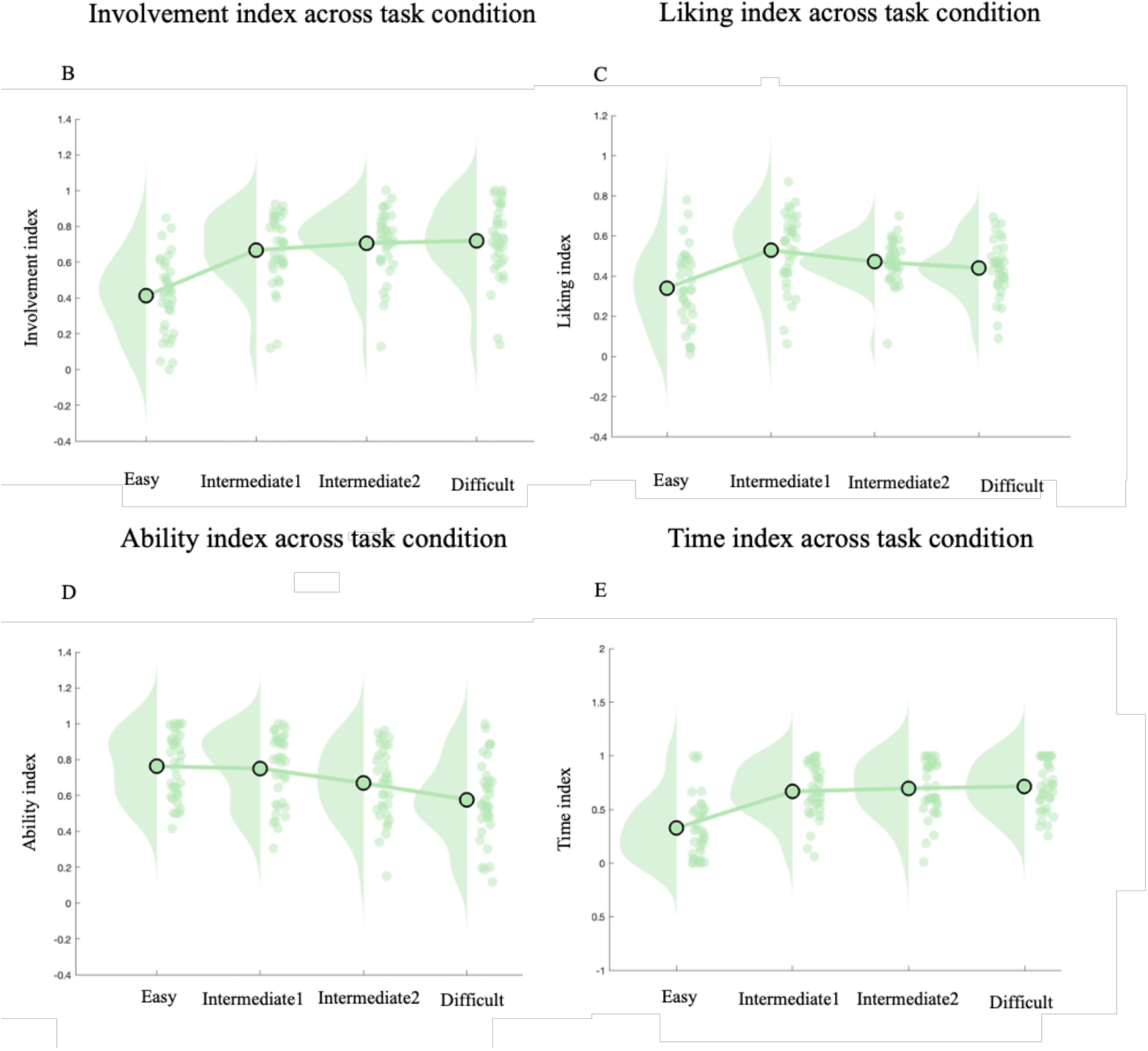
Raincloud plot for A) involvement index, B) liking index, C) ability index, D) time index across task condition. The dots display each participant’s mean score. Circled dots stand for group averages.

### Supplementary Results 2

#### The effects of task related parameters on pupil size and the motivational and behavioural relevance of pupil size only in volunteers below 36 years old

We repeated the analysis performed in sections 3.4. - 3.6. only in volunteers that are below 36 years old. We show that the results qualitatively do not change when volunteers above 36 years old (N=3) are excluded from analysis.

**Supplementary Fig 3.**
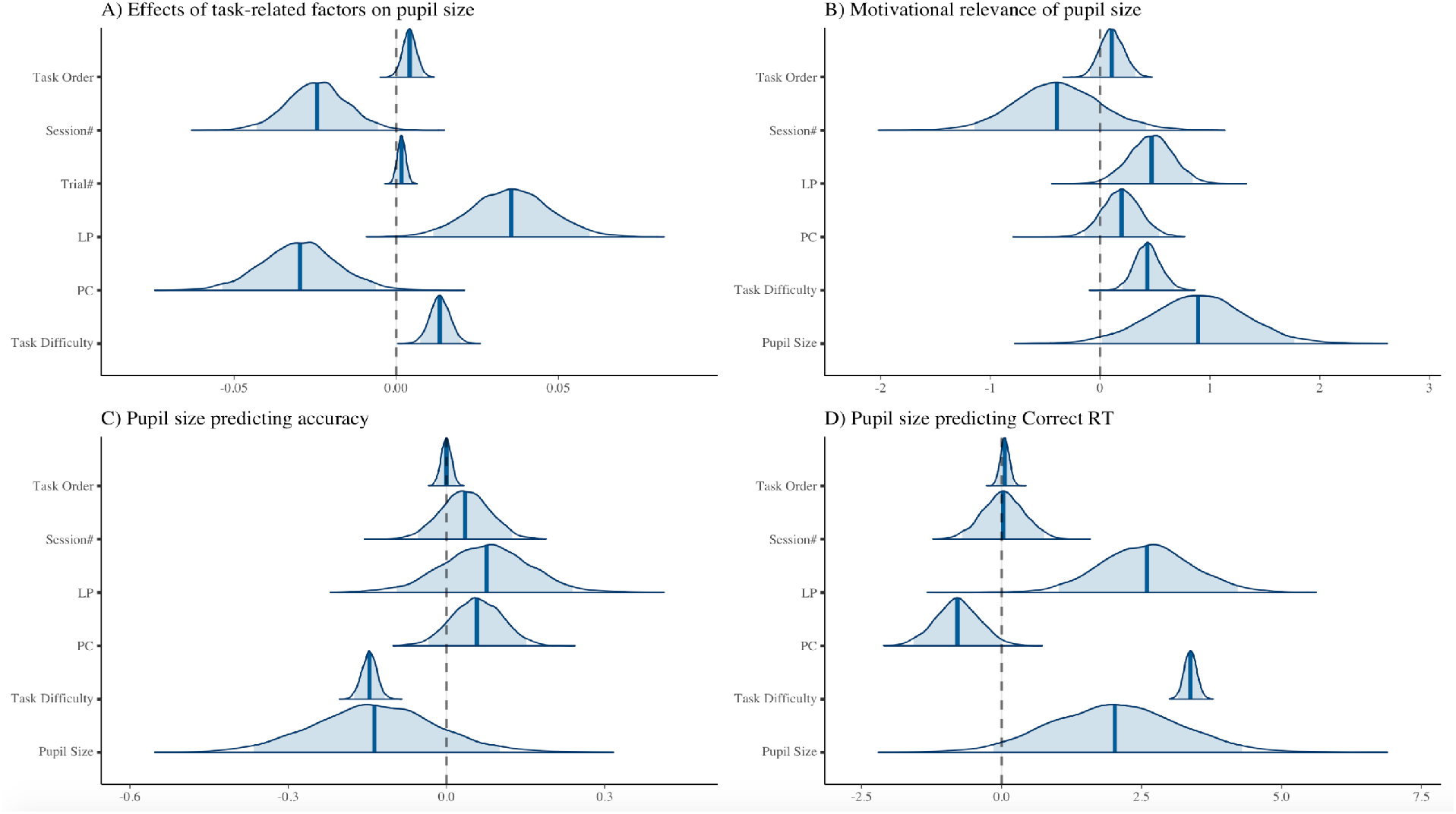
Densities of model parameter estimates. A) Model parameter densities for predicting pupil size. B) Model parameter densities for predicting flow scores. C) Model parameter densities for predicting accuracy. D) Model parameter densities for predicting Correct RTs.

### Supplementary Results 3

#### The effects of task related parameters on pupil size before the cue period

We repeated the analysis performed in sections 3.4. - 3.6. for the pupil size immediately before the onset of the cue (200 ms before cue onset). We show that the effect of trial-by-trial changes in task performance on pupil size qualitatively do not change when the pupil size is defined as prior to cue onset.

**Supplementary Fig 4.**
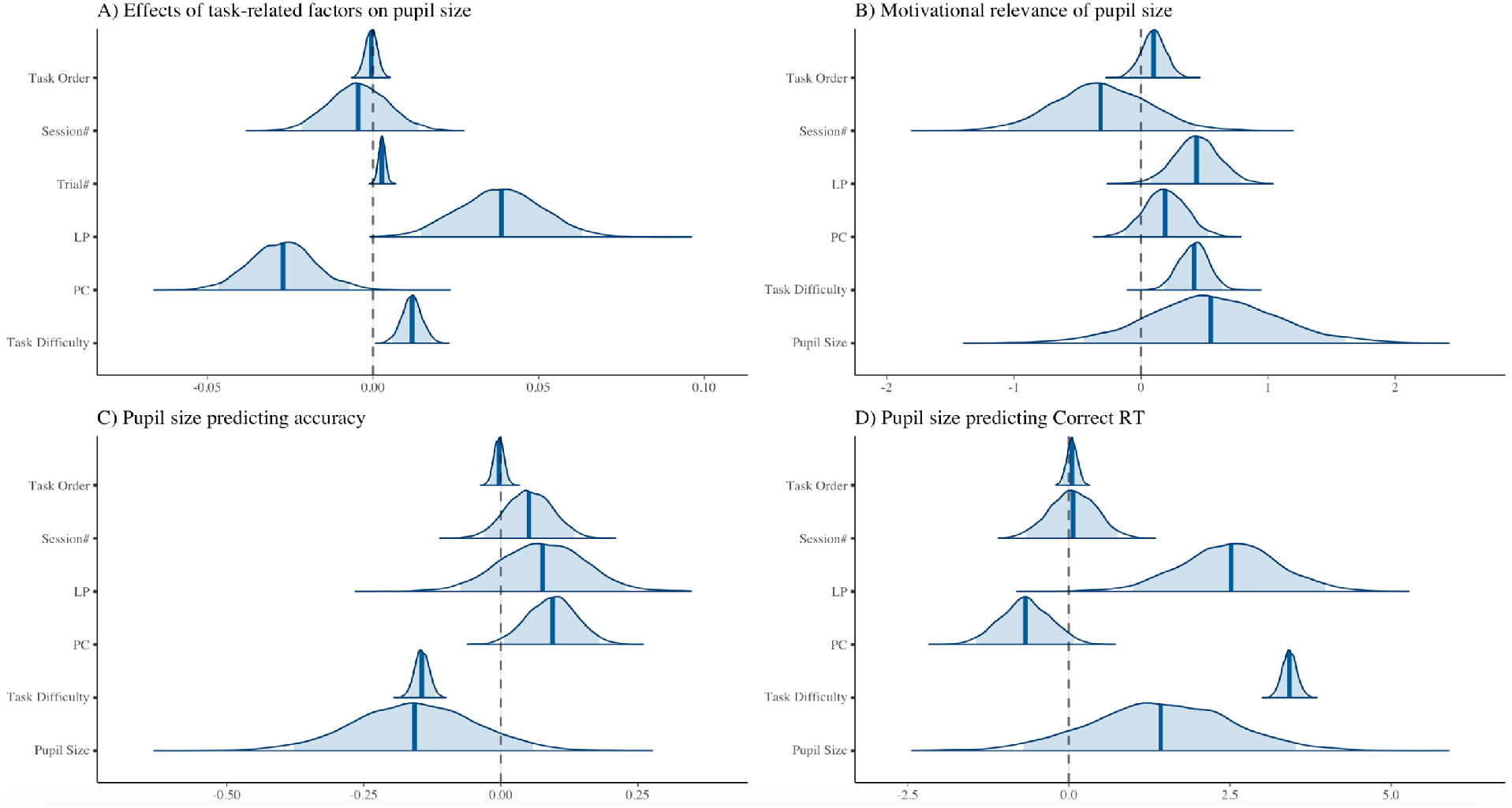
Densities of model parameter estimates. A) Model parameter densities for predicting pupil size. C) Model parameter densities for predicting flow scores. C) Model parameter densities for predicting accuracy. D) Model parameter densities for predicting Correct RTs.

